# Mycobacterial Topoisomerase I Energetically Suffers From C-Terminal Deletions

**DOI:** 10.1101/2024.09.09.612055

**Authors:** Dillon Balthrop, Deepesh Sigdel, Chunfeng Mao, Yuk-Ching Tse-Dinh, Maria Mills

**Affiliations:** Department of Physics and Astronomy, University of Missouri, Columbia, MO 65201, USA; Department of Chemistry and Biochemistry, Florida International University, Miami, FL 33199, USA; Biomolecular Sciences Institute, Florida International University, 11200 SW 8 St, Miami, FL 33199, USA

## Abstract

Type IA topoisomerases relieve torsional stress in DNA by a strand-passage mechanism, using the strain in the DNA to drive relaxation. The topoisomerase IAs of the Mycobacterium genus have distinct C-terminal domains which are crucial for successful strand-passage. We used single-molecule magnetic tweezers to observe supercoil relaxation by wild type *Mycobacterium smegmatis* topoisomerase IA and two C-terminal truncation mutants. We recorded distinct behaviors from each truncation mutant. We calculated the free energy stored in the DNA as it is twisted under force to examine the differences between the proteins. Based on our results, we propose a modified model of the strand-passage cycle.

## Introduction

During various cellular processes torque is exerted on DNA causing torsional strain [1]. This strain can either twist the DNA about its helical axis or contort that axis in three dimensions, causing the DNA to wrap over itself, forming plectonemes. To describe this topology, the parameters twist (Tw) and writhe (Wr) are used. A DNA’s Tw is the number of rotations it makes around its helical axis (Fig. 1A), whereas the number of three-dimensional crossings it makes with itself is called Wr (Fig. 1B). Initially, Tw has lower energy than Wr. As Tw builds up Wr becomes more favorable, causing buckling. Tw and Wr can coexist and interconvert [2] (Fig. 1C), and their sum is called the DNA’s linking number (Lk) (Fig. 1) [3]. When relaxed, DNA of arbitrary contour length L has a linking number Lk_0_. Each change in linking number (ΔLk) from Lk_0_ is a supercoil. Rotating the DNA in the direction opposite its natural right-handed helix decreases the linking number [4] creating negative supercoils. Rotating in the same direction increases Lk [4] creating positive supercoils.

**FIG. 1.**
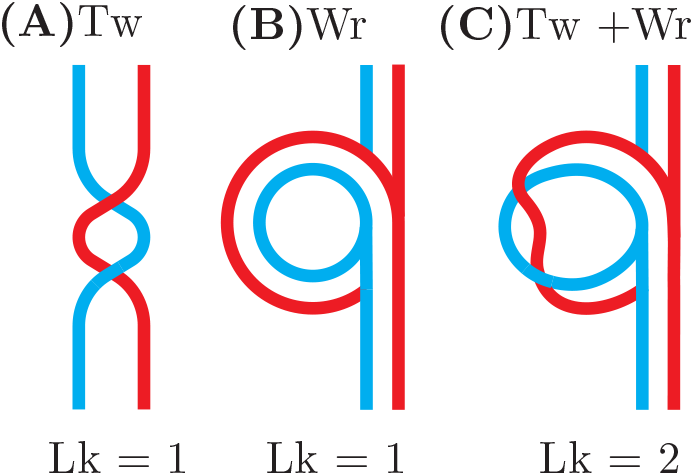
Visual explanation of twist (Tw) and writhe (Wr). Each colored line is one strand of ssDNA. Two strands together form dsDNA. (A) One 360° rotation about the long axis is a twist. One three-dimensional self-crossing of the full DNA backbone is a writhe. (C) Twist and writhe may exist simultaneously.

The accumulation of excess supercoils renders DNA inaccessible, compromising cellular processes [3, 5–7]. To manage DNA topology and relieve torsional stress, a mechanism exists in the form of enzymes called DNA topoisomerases [8–10]. Type IA topoisomerases relax supercoils via a strand-passage mechanism [11–13] (Fig. 2A) in which one strand of the DNA duplex is cleaved (gate strand or G-strand) and the opposing strand is passed through the break (transport strand or T-strand). The broken G-strand is held open by a protein-mediated DNA gate [11]. This process removes supercoils in steps of 1 (ΔLk=1) and does not require an input of energy. Instead, the process is driven by thermal energy and torsional strain in the DNA molecule. Because these enzymes require a single-stranded region to function, they typically relax negative supercoils which are more prone to base pair separation, also called melting [2, 14]. Type IA topoisomerases are ubiquitous but there exists structural diversity between organisms. The N-terminal catalytic core (domains 1-4 or D1-D4) remains highly conserved from the archetypal *E. coli* topoisomerase IA and binds to the G-strand [15–17]. This core forms a ‘padlock’ shape and contains the active site of the protein, which is responsible for reversibly cleaving the G-strand (Fig. 2B). Most type IA topoisomerases have additional C-terminal domains (CTDs), which vary in structure and may bind to the T-strand [18, 19]. The topoisomerases of mycobacteria, such as the disease-causing *Mycobacterium tuberculosis*, have four CTDs (Fig. 2B) which are essential for strand passage [18–20], though their precise dynamics are unclear. The mycobacterial topoisomerase IA CTDs are not present in human topoisomerases, making those domains potential targets for antibiotics [21, 22].

**FIG. 2.**
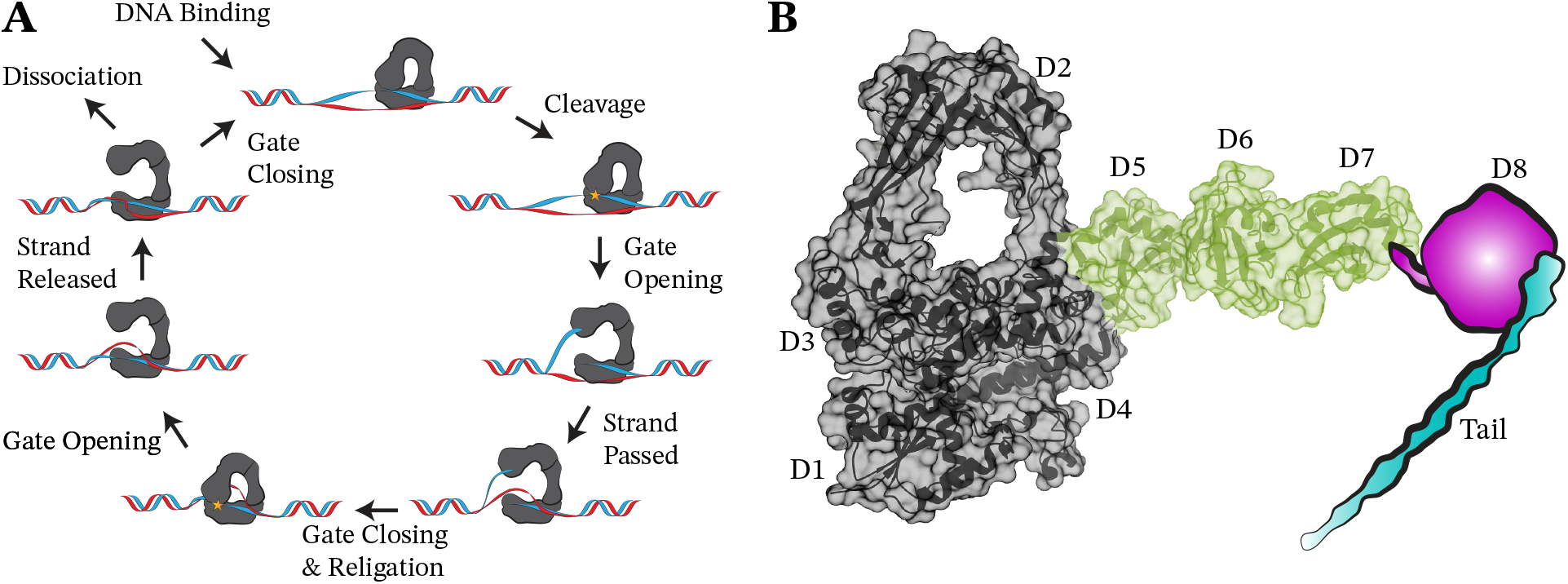
(A) Current proposed strand-passage mechanism for type IA topoisomerases. The C-terminal domains are not shown. (B) Crystal structure of 839t MsmTopI deletant (PDB: 6PCM [19]) with missing domains sketched in. The N-terminal catalytic core is represented in gray. The domains are colored by presence in each mutant. Domains in the 839t mutant are D5-7. The domains present in the 909t mutant are D5-8. Throughout this work, data and schematic coloration are the same.

Previous work has revealed the general role of the of the domains and their impact on supercoil relaxation [18, 19, 23]. *Mycobacterium smegmatis* TopI (MsmTopI), the focus of this study, is a common model for *Mycobacterium tuberculosis* TopI. Mycobacterial TopI has four C-terminal domains (D5-D8) and a 27-residue long, basic, and unstructured tail (Fig. 2B). These CTDs have high affinity for ssDNA binding [18, 19]. Bulk biochemical experiments have shown that sequential deletions of MsmTopI CTDs resulted in exponentially diminishing relaxation activity [19]. Others have also shown that removing basic portions of the CTDs reduces activity [18]. The same truncated mutants used by Cao and colleagues [19] are investigated in this work. The first truncated mutant, 909t MsmTopI (909t), lacks the 27-residue, basic, unstructured tail, and terminates at the 909th residue at the end of D8. The second truncated mutant, 839t MsmTopI (839t), lacks both the tail and D8. Our work aims to closely examine how these deletions affect the relaxation process on a single-molecule scale.

Single-molecule techniques are ideal for probing the details of such protein mechanisms [24–28]. Single-molecule Magnetic Tweezers (MT) are commonly used to analyze protein-nucleic acid interactions due to their ability to record and observe data in real-time, capture multiple signals at once, and the small amount of perturbation being applied directly to the system [29–32]. MT consist of an inverted microscope with a pair of overhanging permanent magnets. The magnets can be rotated clockwise (+) or anti-clockwise (-) to introduce supercoils (SC) to the DNA. The magnets may also move up or down to vary the force applied to the DNA (Supplementary Fig. S1). Overall, we observed distinct differences between all three enzymes. Ultimately, to link the loss in activity seen in bulk experiments and individual characteristics seen in single-molecule experiments, we looked to the DNA’s changing energy as supercoils were relaxed and what possible steps in strand-passage could contribute to the observed behavior.

### Relaxation Analysis

We measured the DNA relaxation activity of MsmTopI by an established MT assay [11, 29, 33]. Functionalized 6 kb DNA tethers were attached on one end to the bottom coverslip of a flow cell and on the other end attached to a paramagnetic bead. The tethers were held at a constant force of 0.45 pN. This force causes slight melting of negatively supercoiled DNA and creates a single stranded binding region for the enzyme. The tethers were negatively supercoiled (-SC) by 30 turns in the presence of 1 nM WT MsmTopI. The z-positions of the beads were recorded in real time as the enzyme relaxed the supercoils. Once the tethers were relaxed, they would then be recoiled, and the process would be repeated. Over the course of a single relaxation event there were typically multiple processive runs interspersed by pauses, or dwells, of varying durations (Fig. 3). The data was then fed to a custom Python algorithm which implemented Student’s T-test to identify the start and end points of each relaxation run and parameters of interest were calculated. The results were plotted as histograms which were then fit with the appropriate distributions to obtain statistical quantities (Fig. 3). Experiments and analysis were performed identically for the mutants. For each enzyme, there was a range of 5 to 30 events per experiment, with a minimum of 10 experiments performed.

**FIG. 3.**
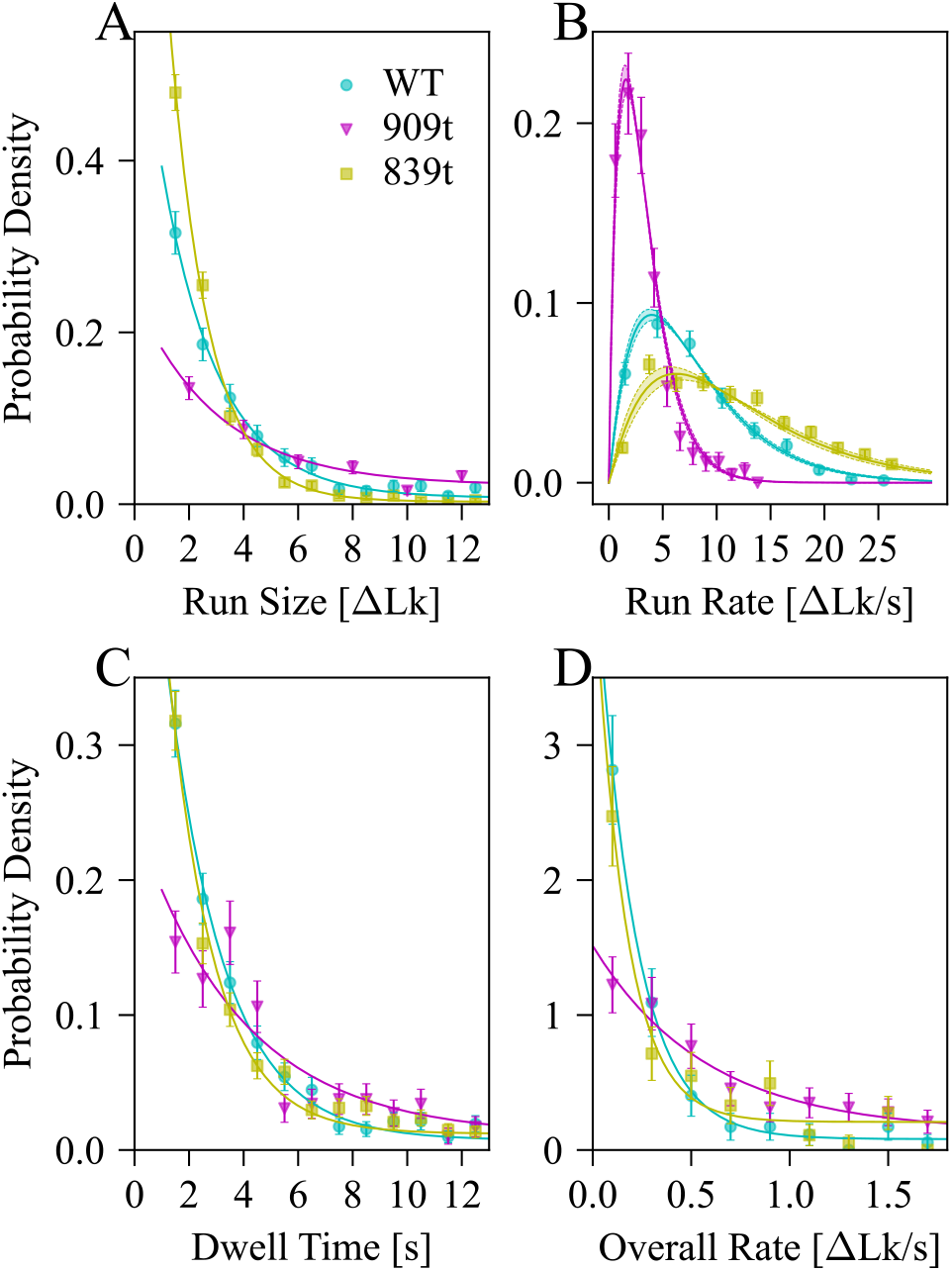
Histograms showing the distributions for different properties that were analyzed. All data was normalized to a probability density. (A) Histogram of run size. (B) Histogram of run rate. (C) Histogram of dwell times. (D) Histogram of overall rate.

While type IA topoisomerases are known to change the DNA linking number by discrete steps of 1 per kinetic cycle [34], they are also known to stochastically complete more cycles in a processive burst or run [32, 33]. In our observations (Table I) the Wild Type (WT) enzyme relaxed 4.7 ΔLk, on average, per run with an average run rate of 7.9 ΔLk/s (Fig. 3A-B). The 909t mutant exhibits a notable preference for longer average run sizes of 8.7 ΔLk, but a slower run rate at 3.3 ΔLk/s. Run sizes for the 909t also spanned a wide range, with the enzyme relaxing up to 25 turns in a single run in some instances. The 839t has a short average run of 3.2 ΔLk, but works at a much faster run rate of 12.2 ΔLk/s. Considerable variance was observed in dwell times between relaxation events (Fig 3C). The WT dwell times averaged 6.59 seconds between runs with frequent instances of dwells exceeding 1 minute. Analysis of dwell positions for the WT revealed a preference for dwelling primarily within the first and last 20% of relaxed turns (Fig. 4). However, there was no correlation between where a dwell occurred and its duration. In fact, the WT maintained relatively consistent pacing regardless of supercoil density. Average dwell times for the 909t mutant were longer at 8.14 seconds and more likely to occur with longer durations as more of the supercoils were removed. The 839t displayed dwell durations shorter than the WT with an average value of 6.14 s but has more dwells overall due to its smaller run sizes. Interestingly, the 839t does show some instances of prolonged dwells near Lk_0_, but the large number of dwells overall renders them statistically insignificant. Increases of dwell times with dwell positions can be contextualized by the overall relaxation rate, which is defined to be the change in Lk per second over the time of a full relaxation event. The WT’s overall rate was determined to be approximately 0.34 ΔLk/s (Table I, Fig. 3D). The 909t overall relaxation rate is 0.67 ΔLk/s. This is despite its exceedingly long dwell times at Lk*≈*Lk_0_. The 839t mutant achieves an overall rate of 0.52 ΔLk/s, surpassing the WT but remaining under the 909t mutant’s rate. Qualitatively speaking, the 909t relaxes more turns without stopping, reducing the impact of its lower run rate by eliminating extra dwells, and slows down dramatically as Lk→Lk_0_. In contrast, the 839t relaxes in rapid and short bursts, but is plagued by many dwells of varying duration which pull its overall rate below the 909t’s.

**TABLE I.** Overview of experimental results. Values are from fits to histograms, and errors are from 2 standard deviations.

| Protein | Run Size [$\Delta\text{Lk}$] | Run Rate [$\Delta\text{Lk/s}$] | Dwell Time [s] | Overall Rate [$\Delta\text{Lk/s}$] |
| --- | --- | --- | --- | --- |
| Wild Type | $4.73 \pm 0.70$, $n_{rs} = 549$ | $7.9 \pm 0.1$, $n_{rr} = 529$ | $6.59 \pm 1.10$, $n_{dw} = 552$ | $0.34 \pm 0.08$, $N = 105$ |
| 909t | $8.74 \pm 1.60$, $n_{rs} = 421$ | $3.3 \pm 0.1$, $n_{rr} = 411$ | $8.14 \pm 3.00$, $n_{dw} = 407$ | $0.67 \pm 0.25$, $N = 159$ |
| 839t | $3.21 \pm 0.80$, $n_{rs} = 1217$ | $12.2 \pm 0.1$, $n_{rr} = 1217$ | $6.14 \pm 0.90$, $n_{dw} = 1217$ | $0.52 \pm 0.14$, $N = 128$ |

**FIG. 4.**
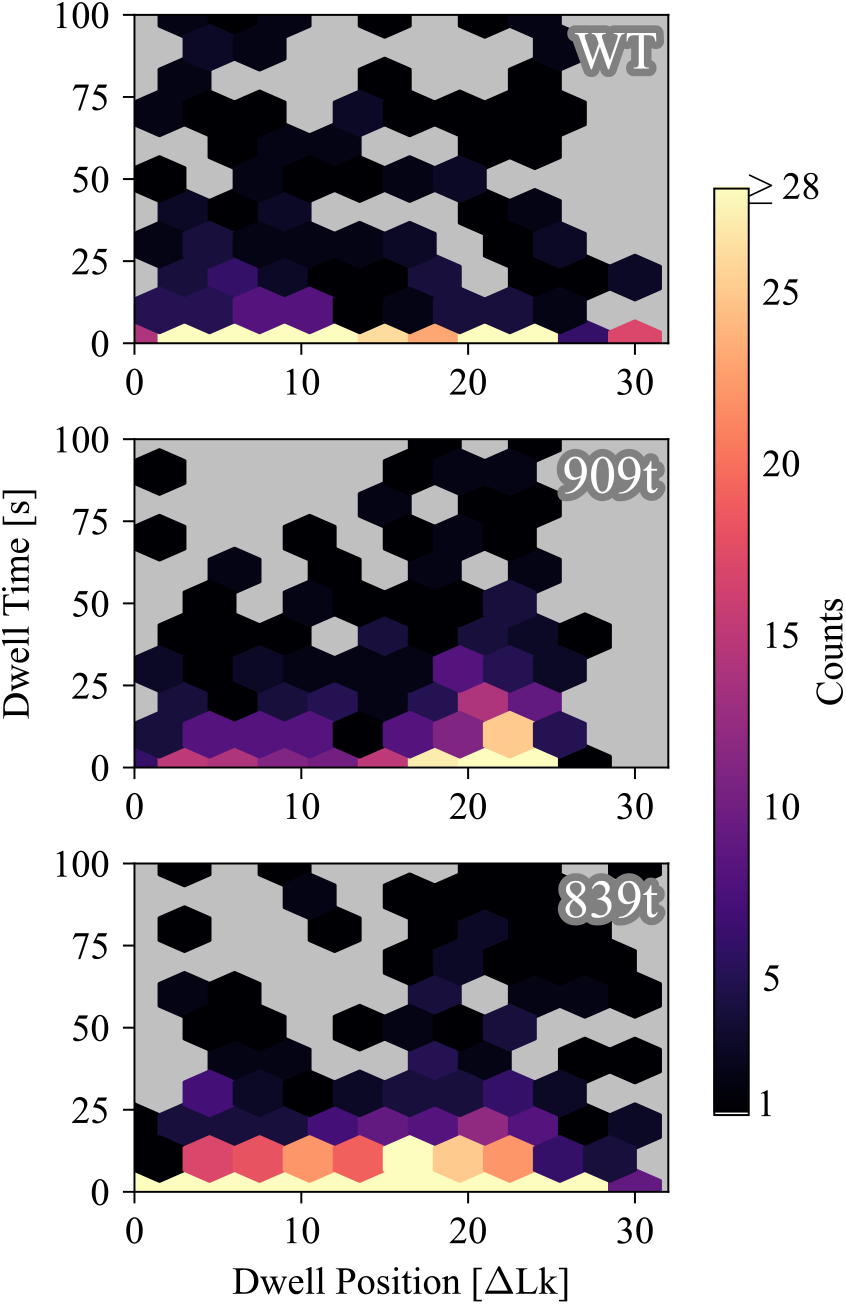
Correlation heat maps for dwell times and dwell positions of each enzyme. Points which are high on the y-axis are longer dwells. Points which occur further on the x-axis represent dwelling after more supercoils were resolved from the substrate.

Given that the 909t was expected to be *∼*1/2x as effective as the WT, and the 839t *∼*1/18-36x based on previous results from bulk assay of relaxation of supercoiled plasmid, our results were initially puzzling. However, the overall rate only encapsulates the rate from start to stop of activity, and not all events ended with the DNA completely relaxed. We analyzed each protein’s stopping Lk (Fig. 5A), defined as the ΔLk from fully relaxed after 30 minutes of no activity on the tether. Unimodal gaussian distributions describe both the WT and 909t, centered at 8 and 10 ΔLk respectively (Fig. 5A). The WT’s distribution was broad, ranging from 0-16 ΔLk remaining, but the 909t’s was tighter, only covering from 5-15 ΔLk remaining. Surprisingly, the 839t displayed a bimodal gaussian distribution with peaks centered at 11 and 4 ΔLk remaining, but cumulatively, it spans the same range as the WT.

**FIG. 5.**
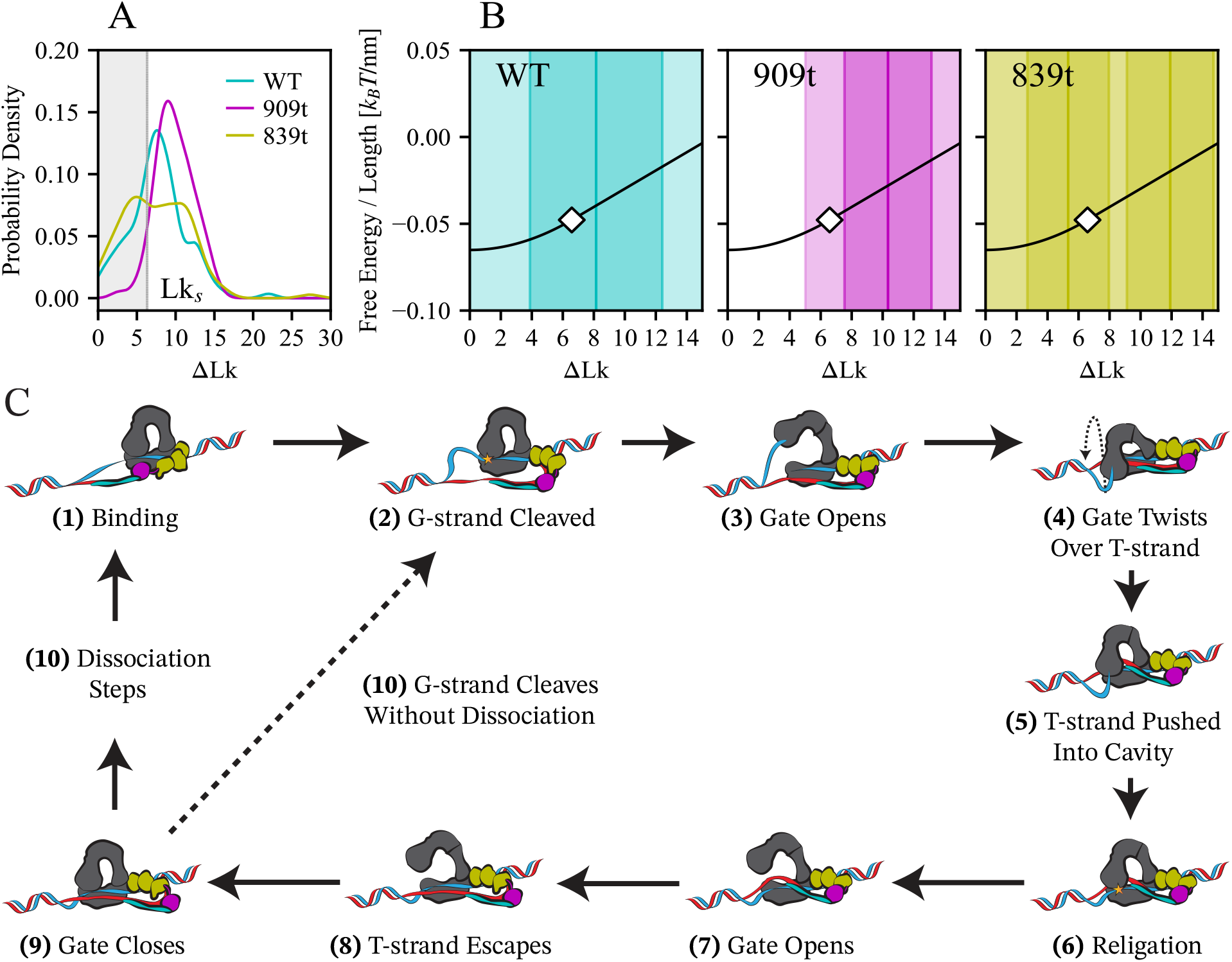
Plots demonstrating the connection between the free energy profile of the DNA matching our experiment conditions, its buckling point, and how much linking number was remaining in the DNA before enzyme activity ceased. (A) Smooth histograms of the enzymes’ recorded stopping positions. The dashed gray line represents the DNA buckling transition Lk_*s*_=6.38 ΔLk. ΔLk in the shaded region have only twist and no writhe, and ΔLk outside the shaded region have both twist and writhe. (B) Overlays of each protein’s distribution from (A) on the free energy profile of the DNA. The solid black curves represent the DNA’s free energy as a function of Lk, with a white diamond marking the DNA buckling transition. The solid vertical lines are the peaks of the Gaussians which fit the data. Regions are shaded dark within the full-width half-maximums of each Gaussian, and lightly shaded for the remaining data. (C) Our proposed strand-passage cycle which implements DNA corralling.

The differences in stopping positions illustrate that the loss of the CTDs affects strand-passage differently depending on how much supercoiling has been removed. In contrast to the WT, the 909t struggles to relax turns near Lk_0_. This suggests that the 27-residue tail plays an important role in supercoil removal as Lk approaches Lk_0_. This could be due to decreased binding to the T-strand and lower energy remaining in the DNA for relaxation. Lysine and arginine residues present in the tail imply strong electrostatic interaction with the T-strand and give credibility to these possibilities. Given that the proteins use the torsional strain in the DNA to perform relaxation, it is likely that the DNA buckling transition plays a role in all these distributions. Further, this energy deficit could be exacerbated in the 839t that has one less C-terminal domain for binding to T-strand. To probe this in detail, we looked to the established model by Marko (2007) [2] which connects the DNA free energy to its level of supercoiling under force.

### Energetic Connection

Relaxed DNA has a bend persistence length (*A*) which is the minimum length of DNA required for it to bend, and a twist persistence length (*C*) which is the length of DNA typically spanned by 1 Tw. *C* is also referred to as twist stiffness. The twist persistence length becomes shorter after supercoiling and the new value is denoted as *P*. Similarly, the twist rigidity of stretched DNA under a force, *f*, is defined to be *c*_*s*_, which governs the amount of twist that can be absorbed into writhe. The presence of some constants makes it convenient to rewrite *C* and *P* as 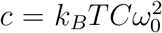 and 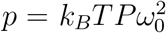 respectively (*k*_*B*_*T* = 4.1 pN*·*nm). It is important to note that various factors, most importantly salt concentrations, can influence the values of *A,c,p* and *c*_*s*_. The supercoiling density, *σ* ≡ ΔLk/Lk_0_, is used in place of linking number for calculation since it is a normalized quantity. There are 3 stages of DNA supercoiling, each with independent expressions for their free energy. First, all ΔLk is added in the form of Tw. The free energy per length of this region is *S*(*σ*; *f*). Once the supercoiling density reaches a critical value, *σ*_*s*_, the DNA buckles, entering the second stage and introducing the coexistence of Tw and Wr. This region’s free energy per length is denoted as *F*(*σ*; *f*). The addition of Wr can be thought of as an exchange of extension for structural stability which keeps the torque constant. As *σ* continues to increase, it approaches another critical value, *σ*_*p*_ and enters the third stage, where all Tw is converted to Wr, and the DNA is all plectonemic. In the case of stages 2 and 3 where the supercoils are negative, considerable DNA melting can occur at forces greater than 1 pN. Because our experiments were done at *f* ≃0.45 pN, our negatively super-coiled DNA is melted enough to have single-stranded regions for the enzymes to bind, but not experience large extension changes from denaturation. Thus, we used the equations for positive supercoiling which are simpler. Simplifying further, we only needed to consider the first and second stages, because once the DNA reaches *σ*_*p*_ it would be coiled all the way to the surface, and there would be no measurable extension change. The equations describing free energy per length of the two regions of interest, respectively, are

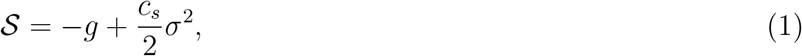

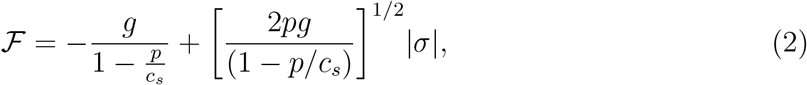

The quantity *g* represents the free energy of stretched, freely swiveling, DNA. To verify values for *A, C*, and *P*, we measured the extension change of initially relaxed DNA tethers as supercoils were added in the absence of protein and compared the results to predictions from the theory. To prevent confusion in data presentation, plots exhibiting these parameters have been converted to linking number (ΔLk) instead of supercoiling density (*σ*) when possible, see Fig. 5. Changing them to use linking number also allows for direct comparison to our experimental extension curves (Supplementary Fig. S2). It is important to note that precision of measured extension is heavily lessened once a tether has relaxed to the buckling transition (Lk_*s*_) due to the flattening of extension measurements for ΔLk smaller than Lk_*s*_. However, it is still possible to tell that a tether has reached that threshold. After determining the parameter values for our system to use in the equations we overlayed the distributions of the proteins’ stopping positions to examine their relationship with the DNA’s energetic profile (Fig. 5A).

The results show the loss of writhe plays a role in the stopping point for the topoisomerases. The peak of the WT’s distribution is *∼*2 ΔLk before Lk_*s*_ (Fig. 5B), and the peak of the 909t’s distribution *∼*4 ΔLk. In fact, the full-width half-maximum (FWHM) of the 909t excludes the buckling transition entirely (Fig. 5B). The same is true for the first peak and FWHM of the 839t, which is further behind the 909t (Fig. 5B). In general, these energy differences highlight that the enzymes’ functions are not just stochastically dependent on the DNA energy, but that the enzyme’s ability to use the energy suffers with structural change. We hypothesize that the most likely way the protein loses access to this energy is through increased mobility of the T-strand during passage caused by loss of the CTDs. This loss of usable energy combined with the results of section A hint at ways the cycle of strand-passage occurs.

It has been shown by Cao and colleagues that the length of available DNA for the enzymes to bind to plays a role in cleavage [19]. Specifically, that on shorter single-stranded segments the CTDs may compete with the WT N-terminal core for binding, resulting in decreased cleavage and, by extension, relaxation rates. Unlike the WT, the 839t mutant showed increased cleavage on shorter single-stranded segments, which explains the second peak observed for the 839t beyond the buckling transition. Given smaller amounts of denaturation as supercoils are removed, it follows that the 839t would have an easier time binding to the regions which remained while the WT would not.

### Modified Strand-Passage Cycle

In general, it is accepted that the purpose of the C-terminus is to aid in passing the T-strand to the cavity, changing the DNA’s linking number [18, 19, 23]. Our results provide insights about the CTDs effects on T-strand passage. In determining possible steps of the process, several concepts were assumed to remain true. First, the CTDs bind to the T-strand and then undergo a conformational change which moves them away from the gate (Fig. 5C 1-5). Second, strand-passage requires multiple attempts from the protein before it succeeds [35]. Third, the protein may stay bound to the substrate after successful strand-passage. Fourth, the gate of the N-terminal core opens as described previously [11]. Fifth, the tail of the WT is bound adjacent to the other CTDs and there is enough room to accommodate it completely. Finally, we assume the CTDs do not enter the cavity.

Under those assumptions, we propose that strand-passage occurs by a DNA corralling method (Fig. 5C). In this model, when the CTDs move away the T-strand is pulled tight, bringing it closer to the gate (2). After or while this occurs, the G-strand is cleaved, and the gate opens (3). As D3 and D2 move up and away from D1, they twist toward the T-strand. This twisting is driven by a mix of thermodynamics and a restoring torque in the cleaved strand. Once D3 has twisted past the T-strand, the gate closes, trapping the T-strand between D1 and D3 (4). Then, D3 and D2 begin to twist back toward D1, pushing, and ultimately locking, the T-strand into the gate (5). The corralling of the T-strand into the gate tightens the CTDs, forcing them to curl in toward the N-terminal core, and creates a tension in the T-strand. The cleaved strand is religated (6) and D3 detaches from the G-strand, allowing the gate to be opened again (7). Seeking to relieve the stress from the curled CTDs, the T-strand exits the cavity and straightens (8), allowing the domains to move away from the N-terminal core again (9). At this point, the protein may either dissociate or complete the cycle again (10). We attempted to construct this cycle such that when CTDs are deleted, the overall cycle stays the same, but its properties could agree with our results. Each deletion of the CTDs effectively changes the enzyme’s length, charge density, and binding affinity. The combined length of the CTDs, including the tail, could reduce the amount of T-strand which is available for strand-passage; possibly providing better control of the amount of supercoils removed. This could be important in vivo because DNA is typically in a supercoiled state due to its compaction. Further, the positive charge of the tail could reduce the negatively charged DNA’s self-repulsion, allowing the strands to be brought closer to each other, aiding strand passage (Fig. 5C 4-5). Others have shown separating the N-and C-terminal domains and complimenting them in a relaxation assay somewhat restores activity, but the N-terminal domain is unable to relax supercoils by itself [23]. This indicates that the electrostatic and structural changes to the T-strand imparted by the CTDs are critical for strand passage and agrees with our results.

We posit the enzyme’s failed attempts at strand-passage appear as decreases in the enzyme’s run rate and increases in dwells, both frequency and duration, within our data. Run rate is being included because single relaxation steps are below our resolution limits; so frequent small dwells during bursts would be missed. This qualitatively explains the decrease in run rate from the WT to the 909t. Without the tail, the 909t may need more attempts at corralling the T-strand before successfully capturing it. One possibility for this behavior is that the tail stabilizes the T-strand’s position relative to the G-strand, and without it the T-strand can’t be manipulated as easily. Another is that the T-strand may not be able to consistently be brought close enough to the gate to be passed. Considering the activity of the 839t, we believe it to most likely be the latter or a combination of both. Provided that the CTDs move away from the gate, losing D8 may force the T-strand to be brought closer to the gate. This decrease in distance may allow the T-strand to pass in fewer attempts than even the WT. However, strand-passage would still be hindered by the lack of structure and maneuverability in the T-strand, resulting in more pauses and shorter bursts of activity. In other words, it is a trade of consistency for speed. One could expect that as more CTDs are removed, the T-strand distance becomes shorter, but control over it becomes lesser.

## Acknowledgements

We would like to thank Keir Neuman and Ian Morgan for sharing Python code to assist with data analysis, Connor Nance, Dayton Simpkins, Pavel Morales, Dylan Weaver, and Kara Balthrop for numerous long and helpful discussions, and Samuel Cheslik and Doniven Hicks for help with flow cell assembly. This work was supported by laboratory start-up funds from the University of Missouri.

