## Supplemental Material for "Mycobacterial Topoisomerase I Energetically Suffers From C-Terminal Deletions"

### Methods

#### MT Measurements

Measurements were performed on a combined magnetic tweezers-TIRF instrument purchased from MadCityLabs. Sample cells used for experiments were prepared as in Mao and Mills [1]. Experiment buffer solution was 50 mM Tris HCl, 100 mM K-Glut, 1 mM Mg-Glut, 0.01% BSA. An average force calibration versus magnet position for 1 μm MyOne magnetic beads (Invitrogen) was obtained using Brownian motion analysis of 6 kb DNA tethers. The coilable DNA molecules were prepared as described by Seol and Neuman [2]. In a typical experiment, a force range of 0.4-0.5 pN was applied to the bead with the magnetic tweezers to keep the DNA extended and away from surface and causing slight base pair separation. Fields of view with multiple usable tethers were first identified. Tethers were considered usable if DNA could be supercoiled by rotating the overhead magnets, and if they fell within a characteristic length between 1.2 – 1.6 $\mu$m. The magnets were then completely raised to avoid adding stress to the tethers while the desired concentration of enzyme was added. Enzymes were purified as described by Cao et al. [3]. Once equilibrated, the force was set back to the prior value. The z position of the bead was tracked over time to monitor the changes in extension of DNA tether. If the tether was relaxed to its original height, more supercoils would be introduced. This relaxation and rewinding would continue until the DNA became unusable; i.e. would not recoil, were nicked, or detached from the surface. All data were collected at 60 Hz at room temperature (21 ± 2 °C). The image tracking was accomplished with a modified LabVIEW program based on the program of Prof. Michael Poirier of Ohio State University.

#### Data Analysis

Experimental data was first loaded into Igor Pro 9 and subjected to preprocessing, where they were decimated by a factor of 2. They were then exported and analyzed using a custom Python program which uses Student's T-test to identify run starts, and stops, in data traces. The base of this program was provided by Dr. Ian Morgan. It then performs linear fits to the detected regions, and we pulled the relevant quantities from the fits. The algorithm implements thresholding to avoid detection of events which are below our resolution limits. Once values were obtained, they were either further analyzed using Python or binned in histograms using Igor Pro 9. If multiple distributions were seen, the data would be put through another custom Python algorithm which implements information criterion to determine the optimal number of Gaussian distributions that fits the data.

#### Energy Analysis

The analysis of the DNA energy utilizes and follows the formulation from Marko [4]. Important to the usability of this framework is the accurate values of $A$, $C$, and $P$ because they depend on experimental conditions like salt types and concentrations. We determined the parameters for our system by taking several DNA extension curves. Because the derived equations allow for the computation of theoretical extension curves, we optimized each parameter in computation until we saw agreement between the theoretical prediction and our experiments. Once the parameter values are known, one can simply follow the formulation to obtain the equations of free energy which govern the three regimes of supercoiling density. As stated in the main text, we may disregard the third regime because, experimentally, the DNA would be fully supercoiled to the surface at that point, rendering no extension change. After the free energy curve was obtained, we converted the stopping positions to the number of turns from fully relaxed to avoid any biasing from possible error in number of turns added at the start. The distributions for each protein were then overlayed onto the free energy curve.

The predicted extension curves (Supplementary Figure S3) were calculated by taking the negative of the partial derivative with respect to force, $-\partial/\partial f$, of $\mathcal{S}$ and $\mathcal{F}$. The derivatives are representative of the DNA's extension per length, a normalized quantity, denoted as $z/L$ where $L$ is the DNA contour length, and are valid over the previous ranges of $\sigma$. The equations are

$$-\frac{\partial \mathcal{S}}{\partial f} = g + f - \frac{c - c_s}{2}\sigma^2, \quad (1)$$

$$-\frac{\partial \mathcal{F}}{\partial f} = \left[\frac{1}{2f\xi} - \frac{\sigma}{4f}\sqrt{\frac{2p}{g\xi}}\right](f+g) + \frac{Ccp}{8Ac_s^2 f^2 \xi^2}\left[\sigma\sqrt{\frac{pg\xi}{2}} - g\right](f-g). \quad (2)$$

The $\xi$ term is defined to be $\xi \equiv 1 - p/c_s$.

### Supplementary Figures

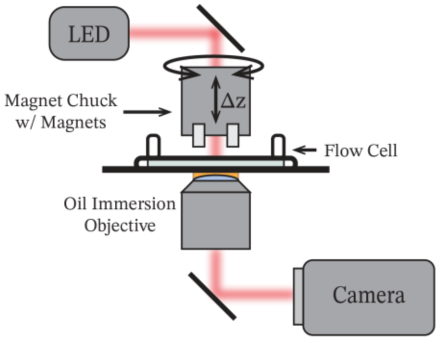


Supplementary Figure S1: Schematic of magnetic tweezers microscope.

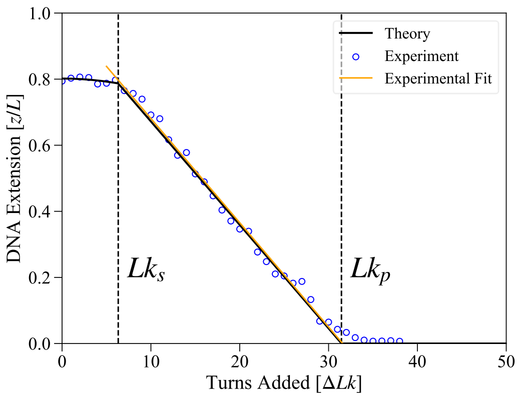


Supplementary Figure S2: Representative hat curve taken from experiment with a linear fit performed on the range between $\mathrm{Lk}_s$ and $\mathrm{Lk}_p$. The y-axis is the DNA extension per length, a normalized quantity where $L$ is the DNA contour length. Used for comparison with the theoretical model employed in the main text to identify proper parameters for the model.

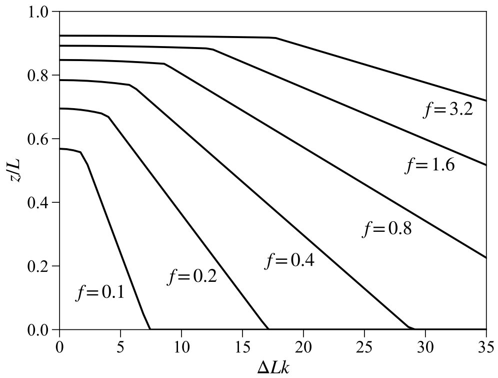


Supplementary Figure S3: Theoretical predictions for DNA hat curves at various forces (in pN). The y-axis is the DNA extension per length, a normalized quantity where $L$ is the DNA contour length. Parameters used for the DNA were determined based on our experimental conditions. Values are heavily salt dependent and were calculated to be $A = 58$ nm, $C = 95$ nm, and $P = 16$ nm.

### Supplemental Material References


[1] C. Mao and M. Mills, Characterization of Human XPD Helicase Activity with Single-Molecule Magnetic Tweezers, Biophysical Journal 123, 260 (2024).

[2] Y. Seol and K. C. Neuman, Magnetic Tweezers for Single-Molecule Manipulation, in Single Molecule Analysis: Methods and Protocols, edited by E. J. G. Peterman and G. J. L. Wuite (Humana Press, Totowa, NJ, 2011), pp. 265–293.

[3] N. Cao, K. Tan, X. Zuo, T. Annamalai, and Y.-C. Tse-Dinh, Mechanistic Insights from Structure of Mycobacterium Smegmatis Topoisomerase I with ssDNA Bound to Both N- and C-Terminal Domains, Nucleic Acids Research 48, 4448 (2020).

[4] J. F. Marko, Torque and Dynamics of Linking Number Relaxation in Stretched Supercoiled DNA, Phys. Rev. E 76, 021926 (2007).
